# A Guide for Population-based Analysis of the Adolescent Brain Cognitive Development (ABCD) Study Baseline Data

**DOI:** 10.1101/2020.02.10.942011

**Authors:** Steven G. Heeringa, Patricia A. Berglund

## Abstract

ABCD is a longitudinal, observational study of U.S. children, ages 9-10 at baseline, recruited at random from the household populations in defined catchment areas for each of 21 study sites. The 21 geographic locations that comprise the ABCD research sites are nationally distributed and generally represent the range of demographic and socio-economic diversity of the U.S. birth cohorts that comprise the ABCD study population. The clustering of participants and the potential for selection bias in study site selection and enrollment are features of the ABCD observational study design that are informative for statistical estimation and inference. Both multi-level modeling and robust survey design-based methods can be used to account for clustering of sampled ABCD children in the 21 study sites. Covariate controls in analytical models and propensity weighting methods that calibrate ABCD weighted distributions to nationally-representative controls from the American Community Survey (ACS) can be employed in analysis to account for known informative sample design features or to attenuate potential demographic and socio-economic selection bias in the national sampling and recruitment of eligible children. This guide will present results of an empirical investigation of the ABCD baseline data that compares the statistical efficiency of multi-level modeling and distribution-free design-based approaches—both weighted and unweighted--to analyses of the ABCD baseline data. Specific recommendations will be provided for researchers on robust, efficient approaches to both descriptive and multivariate analyses of the ABCD baseline data.

## I. Introduction

The Adolescent Brain Cognitive Development Study (ABCD) is a prospective cohort study of a baseline sample of U.S. children born during the period 2006-2008. Eligible children, ages 9-10, were recruited from the household populations in defined catchment areas for each of 21 study sites during the roughly two year period beginning September 2016 and ending in October of 2018. Within study sites, consenting parents and assenting children were primarily recruited through a probability sample of public and private schools augmented to a small extent by special recruitment through summer camp programs and community volunteers. Approximately 9500 eligible, single-born children and 1600 eligible twins completed the ABCD baseline imaging studies and assessments. The sample design and procedures employed in the recruitment of the baseline sample are described in detail in Garavan, et al. (2018).

This methodological paper describes alternative approaches to analysis of the rich array of social, behavioral, environmental, genetic and summary-level neuroimaging data that is collected in the ABCD study. Section 2 will attempt to frame a response to the broad question, why should ABCD analysts be concerned about estimation and inference for the population of U.S. children—does external validity matter? Features of the ABCD design and data that are statistically “informative” and complicate population estimation and inference are the subject of Section 3. Section 4 will attempt to address the specific methodological question, “If inference to the U.S. population is important, what are the appropriate choices of methods for estimating population characteristics and relationships based on the ABCD data?”, describing both model-based and design-based approaches to ABCD estimation and inference. A summary of the general demographic and socio-economic characteristics for the ABCD baseline cohort before any weighting adjustments are applied is presented in Section 5. Section 6 describes the propensity-based weighting adjustment methodology that is used to calibrate the baseline sample cohort to key demographic and socio-economic distributions for U.S. children ages 9 and 10 estimated from the American Community Survey (ACS). Section 7 presents results of an empirical investigation of the ABCD baseline data that compares the statistical efficiency of multi-level modeling and distribution-free design-based approaches—both weighted and unweighted--to analyses of the ABCD baseline data. The paper concludes with specific recommendations for researchers on approaches to both descriptive and multivariate analyses of the ABCD baseline data. Appendices to this paper will contain illustrations of recommended command syntax for analysis of the ABCD data using the major software packages.

## 2. Population orientation to ABCD analysis

As defined in Garavan et al.(2018), the label “population neuroscience “ when applied to observational studies such as ABCD refers to the application of epidemiological research practices including large-scale representative samples to assessments of target populations. It is a study in neuroscience in that it focuses on brain and neurological system development, morphology and function. It is a population study in that observational data are gathered in such a way that they can be used to understand real population distributions and the biological, familial, social and environmental factors that can govern how individuals actually live and grow in today’s society.

From the outset, ABCD’s primary sponsor, the National Institute of Drug Abuse (NIDA) and the ABCD scientific investigators were motivated to develop a baseline sample that reflected the sociodemographic variation present in the U.S. population of 9 and 10 year-old children. ABCD is an observational study sharing many aspects of its longitudinal design with existing population-based survey programs such as the National Longitudinal Study of Adolescent to Adult Health (Add Health, https://www.cpc.unc.edu/projects/addhealth), the Early Childhood Longitudinal Surveys (ECLS, https://nces.ed.gov/ecls/) or the Child Development Supplement (CDS, http://src.isr.umich.edu/src/child-development/home.html) to the Panel Study of Income Dynamics (PSID).

Population representativeness or more correctly, absence of uncorrected selective or informative bias in the subject pool, is important in achieving external validity—the ability to generalize specific results of the study to the world at large. However, even with good, representative samples of populations, failure to measure or control key factors or to recognize important moderating and or mediating relationships can impact external validity of study findings. The ABCD data are observational and although propensity-based methods may be used to control for characteristics of “treated” and “control” participants, in the strictest sense insights gained from the data—even in longitudinal studies such as ABCD—will be associative.

The ABCD baseline recruitment effort worked very hard to maintain a nationally distributed set of controls on the age, sex and race/ethnicity of the children in the study. In year 2, additional monitoring and targeted recruitment were put in place to raise the proportion of children from lower income families. The predominantly probability sampling methodology for recruiting children within each study site was intended to randomize over confounding factors that were not explicitly controlled (or subsequently reflected in the propensity weighting). Nevertheless, school consent and parental consent were strong forces that certainly may have altered the effectiveness of the randomization over these uncontrolled confounders.

The purpose of covariate adjustments in models or the propensity weighting described below in Section 6 is in fact to control specific sources of selection bias and restore unbiasedness to descriptive and analytical estimates of the population characteristics and relationships. For many measures of substantive interest, the success of this effort will never be fully known except in rare cases where comparative national benchmarks exist (e.g. children’s height) from administrative records or very large surveys or population censuses. The effectiveness of weighting adjustments to eliminate bias in population estimates depends of course on the relationship of the substantive variable of interest (e.g. amygdala volume) to the variables that were explicitly used to derive the propensity weights, namely age, sex, race/ethnicity, family type, parental employment status, family size and Census region. These are the types of variables that are available and are identically measured in a national source (American Community Survey) and ABCD. It would have been ideal to have detailed population level data on many other characteristics that may be highly correlated with the ABCD variable of interest (e.g. the child’s parents’ amygdala volume when mom and dad were age 9,10). Only rarely and in large two-phase studies will we ever have population level statistical controls of this nature for a small group such as 9,10 year olds.

“Representative” is a strong adjective to apply to any data set. The accuracy of the descriptor will vary by variable, by subpopulation and by the extent to which the weighting methodology or model covariates capture factors that truly affect the outcome of interest (both in terms of the variables and their functional relationship to the outcome). All forms of statistical estimation and inference make assumptions. No study gets an uncontestable stamp of approval on the unbiasedness of their survey estimates. In both approaches—propensity weighting or covariate adjustment in modeling—it is easy to overlook a selective factor that influences the outcome or modifies the effect of other variables. That is an inherent challenge in population inference from a national study such as ABCD. The position that we take here is that multilevel models that include appropriate statistical controls for demographic and socio-economic factors or propensity weighted estimates of descriptive statistics from the ABCD baseline are in fact publishable estimates for the population of U.S. children so long as authors acknowledge the design and accurately describe the underlying methodology and its assumptions.

## 3. Properties of the ABCD design and data to consider in analysis

This section describes three features of the ABCD design that must be considered in any analysis of the baseline data.

### Clustering and non-independence of observations

Cohort recruitment for the ABCD study design was distinguished by the constraint that eligible children must live within reasonable travel distance (e.g. 50 miles) of a major medical center or research facility where MRI and fMRI imaging could be performed. The geographically-clustered observations on individual children are not independent and the intraclass (“intra-site”) correlations for the many variables must be accounted for to correctly estimate variances of descriptive estimates and analytical model parameters. Correlations among the ABCD observations for individual children are also introduced by other sources of clustering in the ABCD recruitment and measurement protocols: selection of multiple students from schools, multiple children (including twins) recruited from the same family, multiple children imaged on the same MRI scanner.

### Selection bias in site choice and within-site subject enrollment

While the 21 geographic locations that comprise the ABCD research sites are nationally distributed and generally represent the range of demographic and socio-economic diversity of the U.S. birth cohorts that comprise the ABCD study population, in the restricted sense they do not constitute the primary stage of a multi-stage probability sample such as those employed in major population-based epidemiological surveys. To achieve population representativeness for statistical analyses, a mechanism (e.g. modeling site characteristics, assuming pseudo-randomization) is needed to calibrate the broader geographic, demographic and socio-economic characteristics of the set of 21 sites to the larger U.S. population framework (Olsen et al., 2013).

As described in Garavan et al. (2018), within each of the 21 ABCD study sites, a probability sample of the public and private schools was selected as the basis for the recruitment of the majority of eligible children to the ABCD baseline cohort. Although this school-based recruitment approach within each site introduced randomization to the sample of students who could be recruited to ABCD, the process of obtaining school cooperation and then parental consent could selectively impact the final characteristics of the sample that was actually observed. The following sections will describe two approaches, propensity-based weighting and use of appropriate covariate controls in modeling, that aim to address potential selectivity that may have entered the ABCD cohort through the site election or school/parental consent gateways to actual study participation.

### Special twin supplement

A final feature of the ABCD design that deserves attention in the analysis of the baseline cohort data is the special oversample of twin pairs in four of the 21 ABCD sites. Although twins were eligible to be recruited in all sites that used the school-based recruitment sampling methodology, in the four special twin sites supplemental samples of 150-250 twin pairs per site were enrolled in ABCD using samples selected from state registries (Garavan et al., 2018). These special samples of twin pairs can be distinguished in the final baseline cohort of n=11,874 children; however, the study has chosen not to explicitly segregate these twin data from the general population sample of single births and incidental twins recruited through the school-based sampling protocol.

By a default decision of the study team, the propensity-based population weighting methodology described in Section 6 and incorporated in the ABCD Data Exploration and Analysis Portal (DEAP) descriptive estimation does assume a pooled analysis of the general and special twin samples. Section 7 will apply multiple analytic approaches to investigate this assumption that the special twin samples are in fact “exchangeable” with the ABCD general population sample.

## 4. Design-based and model-based approaches to ABCD analysis

Analysts may choose several approaches to estimation and inference that address the challenges posed by the clustering, selection bias and special twin sample properties of the ABCD data.

The first approach is to assume that the multi-stage sample selection for ABCD follows a quasi-probability design and employ design-based methodology similar to that typically used to analyze large probability sample epidemiological surveys such as the U.S. National Health and Nutrition Examination Survey (NHANES). Designed-based analysis will employ population weighting to estimate population statistics and model parameters and non-parametric methods (Taylor Series Linearization, Jackknife, and Bootstrap) to compute robust estimates of standard errors. Any quasi-probability approach to analysis the ABCD data requires a minimum of two things: 1) assignment of cases to ultimate cluster (UC) groupings to account for non-independence of observations; and 2) modeling to derive case-specific analysis weights that account for differential selection factors and permits the observed sample to be “mapped” to the U.S. population of interest (Heeringa, et al., 2017).

As described above, the non-independence of ABCD observations arises through a hierarchical sequence of clustered sample acquisition and measurement stages that include grouping by geographic study site, sampling of schools and clusters of students within schools, familial groupings (primarily twin pairs) of eligible children and even extending to clustered measurement by different MRI scanners within some study sites. Taking the quasi-probability design-based approach, the aggregate impact on variances due to intra-class correlations that arise from these nested sources of clustering can be captured by designating the 21 study sites as the primary stage units (PSUs) or UCs of the sample of children. With case specific weights and PSU codes assigned to each case, weighted estimates and standard errors of population characteristics or parameters in population models can be computed using survey analysis software (e.g. SAS Proc Survey procedures, STATA svy commands:, R Survey Library programs) along with robust TSL, JRR or Bootstrap standard errors and confidence intervals for the weighted estimates (Heeringa, West, Berglund, 2017). A listing of major software system for the analysis of complex sample survey data is provided in Appendix A. Appendix B provides example syntax for ABCD analyses for three of the major survey analysis software packages. A methodology for modeling the population weights for this quasi-probability approach to estimation and inference for ABCD is described in Section 6 below.

ABCD analysts will find that the “design-based” quasi-probability sampling approach is generally the method of choice when the statistics of interest are descriptive estimates of population characteristics such as population totals, means and proportions, quantiles or functions such as a difference of means for two groups (e.g. boys and girls). However, for multivariable analyses designed to explore regression relationships between an outcome and important covariates, multi-level modeling is a second approach that can also be used to explicitly represent the hierarchy of clustering and the associated intra-cluster correlation in the individual observations on ABCD children. The following expression illustrates a mixed effects linear model for ABCD that specifies random intercepts for level 2 clustering of children within families and level 3 clustering of children and families within study sites:

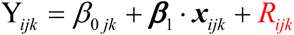

where:

*i* = 1,…, *t* indexes the individual cohort member;

j=1,…, N indexes the cohort member’s family;

k=1,…,21 indexes the ABCD site/imaging center;

***β***_1_ = parameter vector for fixed effects;

*x* = {*x*_1_*_i_*,…, *x_pi_*} = individual level fixed effect covariates;

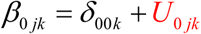 Level 2 random intercept;

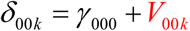 Level 3 random intercept;

for a linear regression model, we assume:

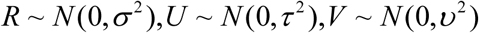

The multi-level mixed model approach to ABCD analysis explicitly disaggregates the total variance of Y into components attributable to each level of the nested hierarchy. While these estimates of variance components may be of interest to analysts, the primary focus of most analyses will be the model estimates of regression coefficients for the Level 1 fixed effects, β.

The multi-level model with random intercepts for families (Level 2) nested with sites (Level 3) will capture the impact of twin-pair and other sibling correlations on variances/covariances of Level 1 (individual) fixed effect estimates. The ABCD Data Exploration and Analysis Portal (DEAP) incorporates this three-level hierarchical specification and uses the R system GAMM4 generalized additive mixed model package as the default for regression modeling of the ABCD data. With added sophistication, analysts can extend the multi-level model, incorporating both random intercepts and random slopes (for site or family covariates) and even cross-level interactions (West, et al. 2014).

Standard applications of the multi-level modeling approach do not incorporate weighting for sample selection or differential nonresponse. Instead, selective characteristics of the observed sample that have a direct effect on the outcome of interest or moderate the effect of other independent variables are included as covariate controls in the estimation of model parameters. In cases such as survey data with population weights that account for unequal probabilities of sample inclusion or differential nonresponse, some authors have suggested that the weight variable itself be included as a covariate in the model specification as further protection against sample selectivity that may not be captured in the observed covariate controls (Rubin, 1996).

Over the past two decades there has been increasing attention paid to weighted estimation of multi-level models for observational data both in the context of inverse probability weighting (IPW) for exposure probability in estimates of treatment effects (Austin and Stuart, 2015) or in analysis of multi-level data (Pfeffermann, et al., 1998). Weighted estimation of multi-level models requires a disaggregation and scaling of the individual propensity or population weights for each level of the model (Rabe-Hesketh and Skrondal, 2006). The ABCD DEAP does incorporate a population weight in estimation of univariate descriptive statistics; however, no IPW or population weighting is currently incorporated in DEAP GAMM4 regression modeling. This paper does not evaluate population weighting for multi-level modeling of ABCD data but work is ongoing on this topic and will be reported in a future update to this guide on ABCD analysis methods.

## 5. Properties of the ABCD Baseline Sample Cohort in Comparison to ACS

The American Community Survey (ACS), a large probability sample survey of U.S. households conducted annually by the U.S. Bureau of Census, provides a benchmark for selected demographic and socio-economic characteristics of U.S. children ages 9,10. The 2011-2015 ACS Public Use Microsample (PUMS) file provides data on over 8,000,000 sample U.S. households. Included in this five-year national sample of households are 376,370 individual observations for children age 9, 10 and their households.

Table 1 compares unweighted demographic distributions for the ABCD baseline cohort to nationally representative ACS estimates for the U.S. population of 9, 10 year olds. With some minor differences, the unweighted distributions for the ABCD baseline sample closely match the ACS-based national estimates for demographic characteristics including age, sex, and household size. This outcome can be attributed in large part to three factors: 1) the inherent demographic diversity of the a=21 study sites; 2) stratification (by race/ethnicity) in the probability sampling of schools within sites; and 3) demographic controls employed in the recruitment by site teams. Likewise, the unweighted percentages of ABCD children for the most prevalent race/ethnicity categories are an approximate match to the ACS estimates for U.S. children age 9 and 10. Collectively, children of Asian, American Indian/Alaska Native (AIAN) and Native Hawaiian/Pacific Islander (NHPI) ancestry are under-represented in the unweighted ABCD data (3.2%) compared to ACS national estimates (5.9%). This outcome, which primarily affects ABCD’s sample of Asian children, may be due in part to differences in how the parent/caregiver of the child reports multiple race/ethnicity ancestry in the ABCD and the ACS.

**Table 1:** ABCD Baseline Cohort Demographic and Socio-Economic Characteristics (Unweighted).

| Characteristic | Category | ABCD<br>n=11,874 | ACS 2011-2015 |  |
| --- | --- | --- | --- | --- |
| | | % | $\hat{N}$ | % |
| Population | Total | 100.0 | 8,211,605 | 100.0 |
| Age | 9 | 52.3 | 4,074,807 | 49.6 |
|  | 10 | 47.7 | 4,136,798 | 50.4 |
| Sex | Male | 52.2 | 4,205,925 | 51.2 |
|  | Female | 47.8 | 4,005,860 | 48.8 |
| Race/Ethnicity | NH White | 52.2 | 4,305,552 | 52.4 |
|  | NH Black | 15.1 | 1,101,297 | 13.4 |
|  | Hispanic | 20.4 | 1,973,827 | 24.0 |
|  | Asian, AIAN,<br>NHPI | 3.2 | 487,673 | 5.9 |
|  | Multiple | 9.2 | 343,256 | 4.2 |
| Family Income | <\$25K | 16.1 | 1,762,415 | 21.5 |
| | \$25K-\$49K | 15.1 | 1,784,747 | 21.7 |
| | \$50K-\$74K | 14.0 | 1,397,641 | 17.0 |
| | \$75K-\$99K | 14.1 | 1,023,127 | 12.5 |
| | \$100K-\$199K | 29.5 | 1,685,036 | 20.5 |
| | \$200K + | 11.2 | 558,639 | 6.8 |
| Family Type | Married Parents | 73.4 | 5,426,131 | 66.1 |
|  | Other Family Type | 26.6 | 2,785,474 | 33.9 |
| Parent Employment | Married, 2 in LF | 50.2 | 3,353,572 | 40.8 |
|  | Married, 1 in LF | 21.9 | 1,949,288 | 23.7 |
|  | Married, O in LF | 1.3 | 156,807 | 1.9 |
|  | Single, in LF | 21.1 | 2,174,365 | 26.5 |
|  | Single, Not in LF | 5.4 | 577,573 | 7.0 |
| Region | Northeast | 16.9 | 1,336,183 | 16.3 |
|  | Midwest | 20.4 | 1,775,723 | 21.6 |
|  | South | 28.3 | 3,117,158 | 38.0 |
|  | West | 34.4 | 1,982,541 | 24.1 |
| Household size | 2-3 | 17.3 | 1,522,216 | 18.5 |
|  | 4 | 33.5 | 2,751,942 | 33.5 |
|  | 5 | 24.9 | 2,085,666 | 25.4 |
|  | 6 | 14.0 | 1,025,285 | 12.5 |
|  | 7+ | 10.3 | 826,496 | 10.1 |

Achieving a balanced nationally representative distribution of children from different socio-economic backgrounds in the baseline sample recruitment presented a major challenge for the ABCD study sites. The 21 ABCD study site populations cover approximately 20% of all U.S. population and certainly include families and children from a wide range of socio-economic backgrounds. ABCD study sites must be located near major research universities or other urban medical imaging centers. Costs of living and therefore basic income requirements for eligible families in the study sites are higher on average than those for the full population of eligible U.S. households. (ABCD is planning to develop alternative measures of ABCD family income that account for local costs of living factors or express a given family income as a percentile of the distribution of incomes for all eligible families in the same site catchment area). In selecting the stratified sample for school-based recruitment of eligible children, the percent of school students who were eligible for Federally-subsidized lunch programs was used as an aggregate measure of the SES level of the student body; however, this school level stratifier was not always a very strong predictor of the SES status of individual ABCD eligible children. The challenge to achieving SES diversity and representative balance in the ABCD baseline cohort originates not only in the population or the limited ability to efficiently stratify and screen on SES characteristics in the sample selection. Parental consent rates and the ability for parent/child pairs to attend the one-day baseline imaging and assessment session are likely to increase with the financial well-being of the family.

The unweighted distribution of reported annual family incomes for the ABCD baseline cohort differs from the nationally representative ACS estimates for the U.S. population of 9, 10 year olds. In nominal dollars, the family incomes of the ABCD children are higher on average than ACS estimates for the comparable U.S. population. Approximately 40.7% of ABCD children are from families with an annual incomes of $100,000 or more compared to an ACS estimate of 27.3% for all U.S. children ages 9,10. As shown in Table 1, ABCD children are also more likely to live in families with two parents who are working—ABCD (50.6%) vs. ACS (40.8%).

The predominantly metropolitan locations for ABCD imaging centers naturally imparted an urban bias to the ABCD baseline cohort recruitment. This bias potential was partially offset at the design stage by defining relatively large catchment areas (e.g. 50 mile radius of study medical centers) for each of the 21 study sites. With few exceptions, the geographic catchment areas extended outward from the more urban centers to include school districts that served suburbs, smaller towns and rural or agricultural areas. Based on 2014 school enrollment data for the 21 study sites, approximately 9% of students in the ABCD catchment areas attended schools that the National Center for Education Statistics (NCES) classifies as “rural”. NCES urban/rural classification coding was used in school sample selection to stratify and enhance the recruitment of students who attended rural schools. Through stratified oversampling of rural schools for ABCD recruitment, the expected proportion of ABCD students enrolled in a rural district rose slightly to 12.5%. This compares to NCES national figures in which 17.5% of ABCD eligible children attend schools in rural districts. In terms of geographic representation of the U.S. household population, ABCD children residing in the Northeast and Midwest Census Regions are represented in approximately the correct percentages; however, overrepresentation of children in the Census West (ABCD 34.3%; ACS 24.1%) is offset by underrepresentation from the large Census South Region.

The ABCD Passive Data Work Group is currently in the process of acquiring external data on school, community and environmental characteristics that can be linked to individual child data and used to analyze the role that these contextual effects may have on the current status and development trajectories of the children in the ABCD baseline cohort. At this stage, the propensity-based population weighting methodology described in the next section does not incorporate calibration based on detailed characteristics of children’s residences, schools or communities.

## 6. Weighting the ABCD Sample to ACS Population Controls

Section 4 above explored the properties of both design-based and multi-level model-based approaches to the analysis of the many forms of data that are collected in ABCD. As noted in that discussion, there is little argument that design-based approaches, common in the analysis of observational data for probability samples, are robust and generally efficient for descriptive estimation of population statistics (e.g. means, proportions, quantiles) and confidence intervals. To that end, a propensity-based population weight has been developed to enable ABCD analysts to apply design-based estimation methods and software for estimating descriptive population characteristics.

In the standard probability sampling framework, the construction of analysis weights for individual observations is organized in three steps: 1) an initial baseline weight to account for planned differences in the sample inclusion probabilities for members of the target population; 2) a model-based adjustment to account for nonobservation/nonresponse in the selected sample; and 3) a final poststratification or calibration of the individual weights to population characteristics known from an external source. Placing ABCD in this framework raises two high hurdles to this standard calculation sequence for population weights. First, ABCD study sites were chosen for their relevant scientific expertise and imaging capabilities and not by a rigorous probability sampling design that assigned each site a known, non-zero inclusion probability. Second, within sites, a probability sample of schools was selected as the primary basis for recruiting eligible children; however, given the available data it is difficult to model the propensities that individual schools and parents (of children in consenting schools) would consent to cooperate in the study recruitment.

Elliott and Valliant (2017) describe two approaches that researchers may take to clearing these two hurdles. Both approaches treat the multi-stage recruitment as a quasi probability sampling process. The first approach takes the hurdles straight on by using detailed modeling to assign quasi-probabilities of sample inclusion (Step 1) and conditional probabilities of study participation (Step 2) to each individual participant. The estimated probabilities are then used to create and assign initial weights which are in turn calibrated to external population controls (Step 3). To create weights for population estimation and inference, ABCD has used a second weighting method that is closely related to the inverse propensity score weighting methodology (IPSW) that is employed to reduce confounding and estimate average treatment effects (ATE) from observational data (Austin and Stuart, 2015). The propensity score method for constructing population weights for an observational data set such as ABCD relies on access to an external, highly accurate and precise “benchmark” data source as a representative gold standard for the population of interest.

As described in Section 4 above, The American Community Survey (ACS), a large probability sample survey of U.S. households conducted annually by the U.S. Bureau of Census, provides such a benchmark for selected demographic and socio-economic characteristics of U.S. children ages 9,10. The first step in benchmarking the ABCD baseline sample weights to population estimates from this large ACS sample required identification of a key set of demographic and socio-economic variables for the children and their households that are measured in common in both in the ABCD assessments and in the ACS household interviews. For the ABCD eligible children, the common variables included:

- age in years
- sex
- race/ethnicity

For the child’s household, additional SES variables included:

- family income
- family type (married parents, single parent)
- household size
- parents’ work force status (modifying family type effect by parent employment status)
- Census Region

Additional candidate variables such as maternal education level were considered for the weight development but were not included due to the fact that ABCD and ACS survey questions were not consistent or one/both of the sources had high rates of item missing data for the variable in question. Once the final set of 8 variables was selected, small amounts of item missing data in both the ACS and ABCD measurements were singly imputed using a chained equations algorithm that is available in the R system MICE package. The data sets containing the common variables from the ABCD (n=11,184) and ACS benchmark (n=376,370) were then concatenated into a single file with each source file distinguished by a binary code (1=ABCD data, 0=ACS data).

Next, a multiple logistic regression model of the following form was fit to the concatenated ACS and ABCD data.

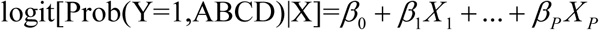

The model is linear in the logit of the probability that the case belongs to the ABCD data. In estimating the parameters of this model, each case in the concatenated file receives a frequency weight. ACS cases are assigned their population weights which in aggregate sum to an average estimate of the U.S. population of children age 9, 10 for the period 2011-2015. ABCD cases are assigned a unit weight (1.0). Applying the frequency weights in the estimation of the model will ensure that the corresponding population propensities for the ABCD sample cases reflect the base population fraction of f=11,874/8,211,605∼ 0.00145 as well as adjustments for the individual covariate factors in the model. As shown below, this base population fraction will correspond to an average population weight for ABCD cases of approximately *W̄* = (0.000145)^-1^ ∼690, with final individual weights varying about this average according to the degree to which the demographic and socio-economic characteristics of the individual case s are underrepresented/overrepresented in ABCD when compared to the ACS benchmark distributions.

The resulting weighted pseudo-likelihood for the binary indicator of ABCD sample membership takes the form:

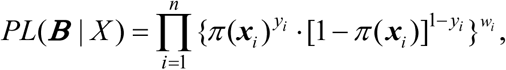

where : w*_i_* = weight assigned to case i in the concatenated ABCD, ACS data set;

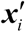 = a column vector of the *p* + 1 design matrix elements for case *i* = [1*x*_1,*i*_…*x_p_*_,*i*_]′;

*π*(***x****_i_*) = the propensity that Y=1 (ABCD) conditional on covariates, ***x****_i_*.

Pseudo maximum likelihood estimates of the parameters are obtained through iterative (Newton-Raphson method) solution of the score equations:

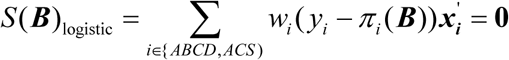

Table 2 provides the estimates of the model parameters and the corresponding adjusted odds ratios for the logistic regression model. Inspecting the beta coefficients and the corresponding adjusted odds ratios, the parameter estimates of the multivariate model confirm the patterns seen in the comparisons of the ABCD and ACS univariate distributions for demographic and socio-economic variables (Table 1). Relative to the ACS population benchmark, the recruited ABCD baseline children have greater odds of belonging to families with two married parents and higher family income. By design, the race/ethnicity composition for major classes (White, Black, Hispanic) matches the ACS target s fairly closely with children of Asian ancestry being underrepresented and children with self-reported Other (AIAN, NHPI, Multiple) race /ethnicity being overrepresented relative to the U.S. population of 9 and 10 year olds.

**Table 2:**
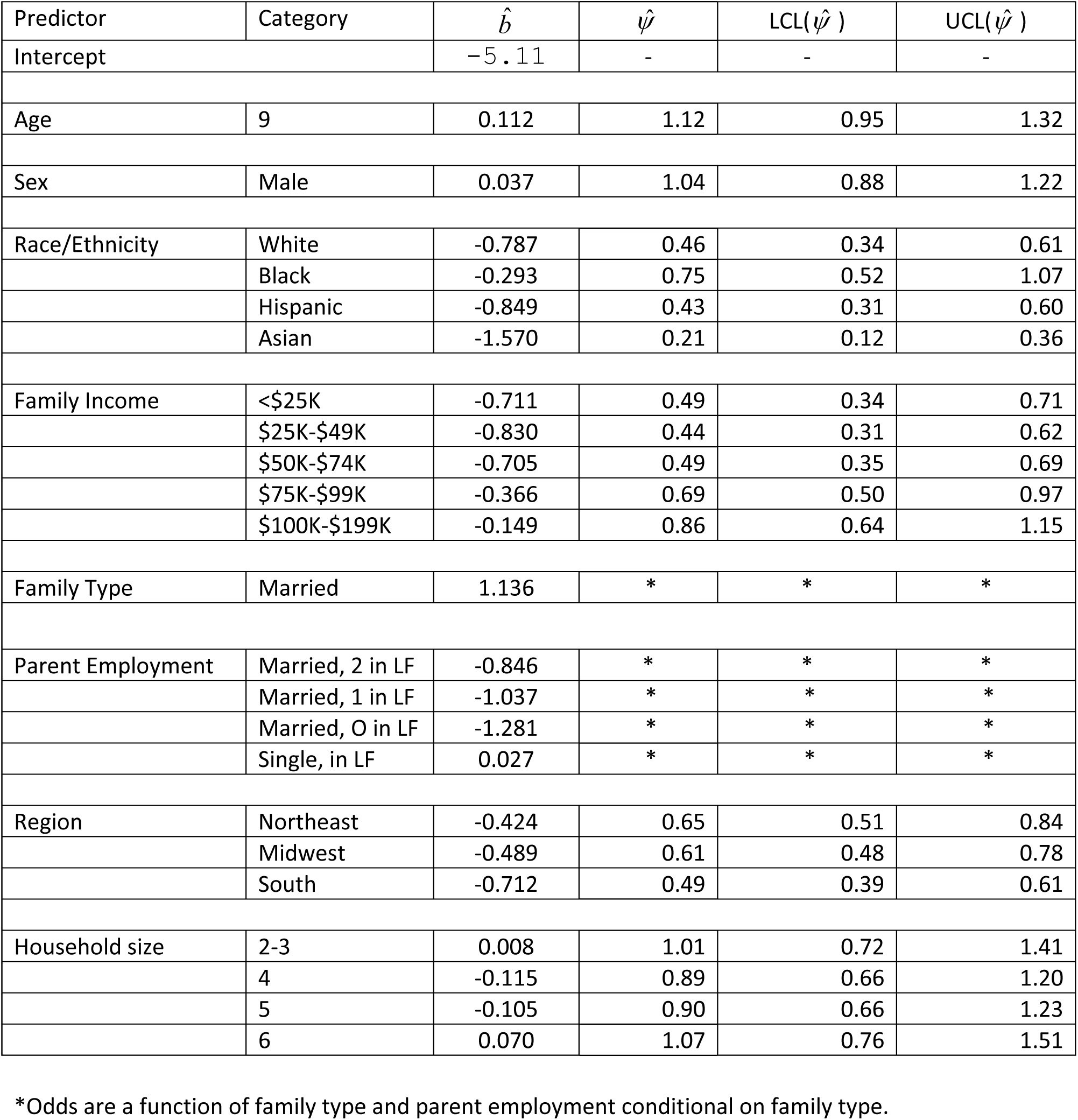
Logistic Regression Model for ABCD Sample Propensity Scores.

Based on the parameter estimates, the predicted propensity that a case belongs to the ABCD data set is obtained using the inverse logit transform:

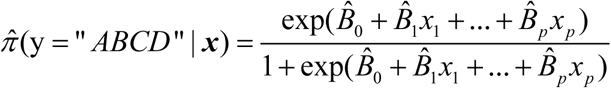

The initial population weight values for each ABCD case are obtained by taking the reciprocal of the predicted propensity for the case.

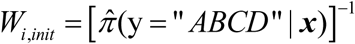

As noted above, the average of the propensity-based population weights for the n=11,874 ABCD baseline cases should be approximately *W̄* = (0.000145)^-1^ ∼690. The actual average of the initial weights was 688.2 (sd=380.22). An equivalent check on the derivation of the propensity-based population weights is to compare the sum of the weights to the benchmark total for the population—the sum of the initial weights assigned to the baseline cohort cases should approximate the population control total estimated from the ACS benchmark which is 8,211,605. For the initial ABCD baseline weights, rounded to two decimal places, the approximation does in fact hold:

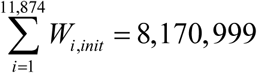

To illustrate the actual weight calculation, consider the initial ABCD baseline population weight value for a 9 year old African-American girl from New England who lives in a family of 4 with two working parents and a family income of $100K-$199K per year:

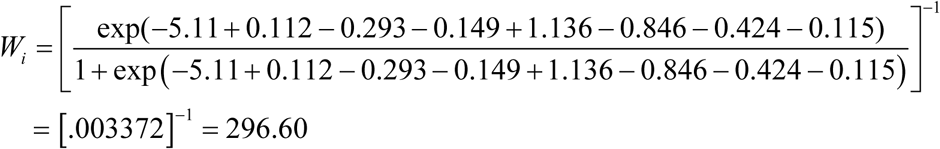

Due to small predicted propensities, some ABCD cases with demographic and SES characteristics that are highly underrepresented when compared to the ACS benchmark population will have extremely large initial weights. Under the estimated model, the child with the lowest propensity of inclusion in the ABCD baseline cohort would be a 10 year old girl of Asian ancestry residing in the South in a 4 person family with two parents who are not working and $25k-$49K total annual income:

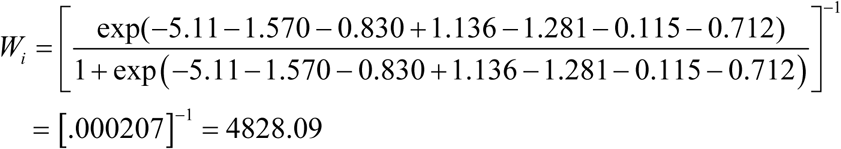

To minimize the impact of the most extreme weights on the variances of descriptive estimates, the extremes of the initial weights were trimmed (“Windsorized”) at the 2% and 98% quantiles of the distribution (Heeringa, et al. 2017). Weights for cases with weight values less than 188 were increased to that 2%-tile value and those with weights greater than 1653 had their weights decreased to 1653 (the 98^th^ percentile).

Following the step of trimming the extremes of the weight distribution, the R Rake iterative proportional fitting algorithm was used to “rake” the trimmed initial weights to exact ACS population counts for the marginal categories of: age (9,10), sex(female, male), and race/ethnicity (Hispanic, Black, White, Asian and all Other persons)—see Table 3. Figure 1 is a histogram display of the frequency distribution of the final population weights for the ABCD baseline children. Figure 2 provides a boxplot comparison of the distribution of weights separately for boys and girls. The relative similarity of these two distributions is evidence of good population balance on the sex of the ABCD baseline participants. The Figure 3 boxplots of weights by family income category show a very different pattern. Compared to the national population, children from families with lower incomes are underrepresented and the population weights for children in these lower income categories have higher average values and a greater variance than the weights for the children from higher income families.

**Figure 1:**
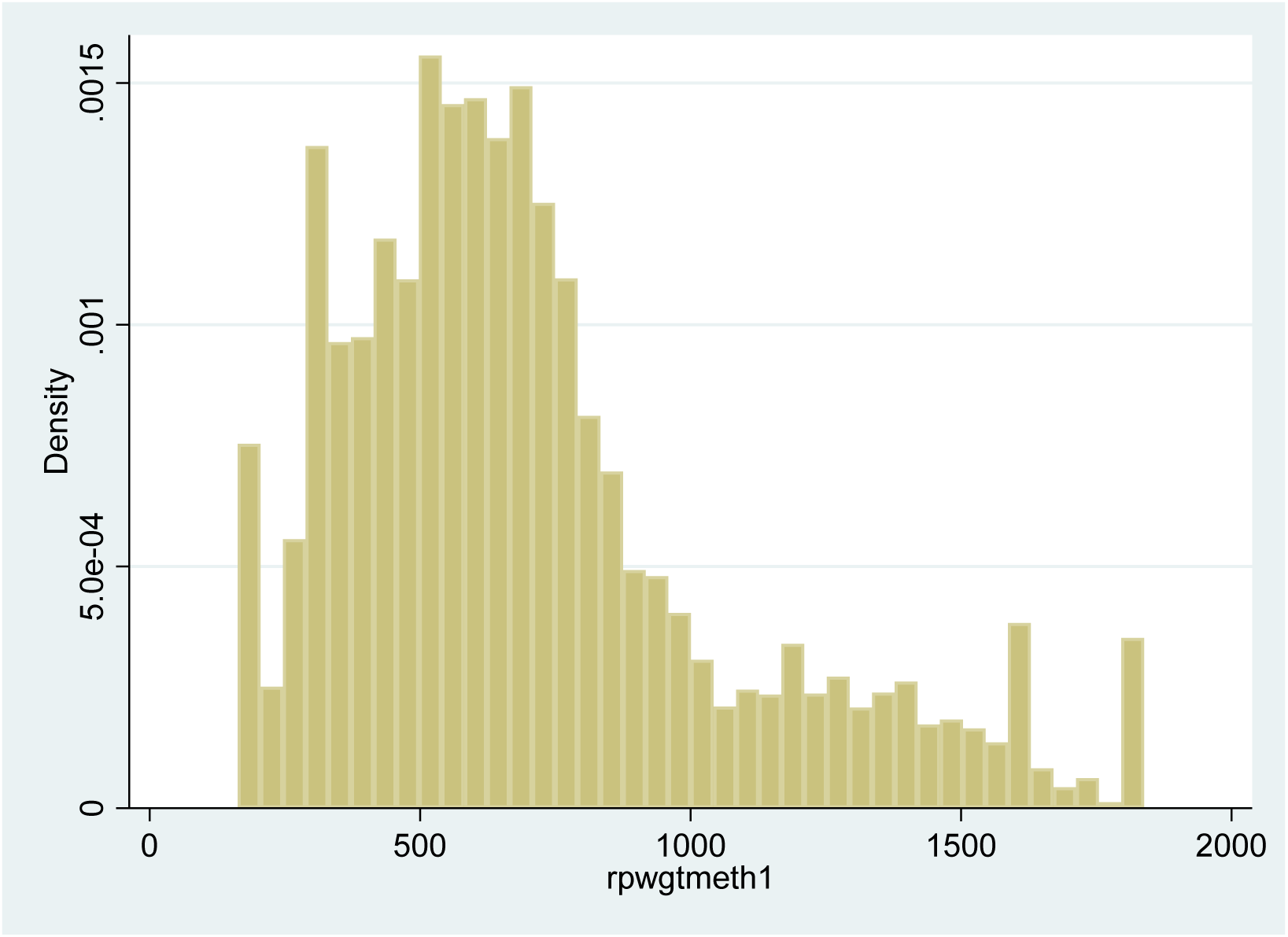
Distribution of ABCD Baseline Population Weights.

**Figure 2:**
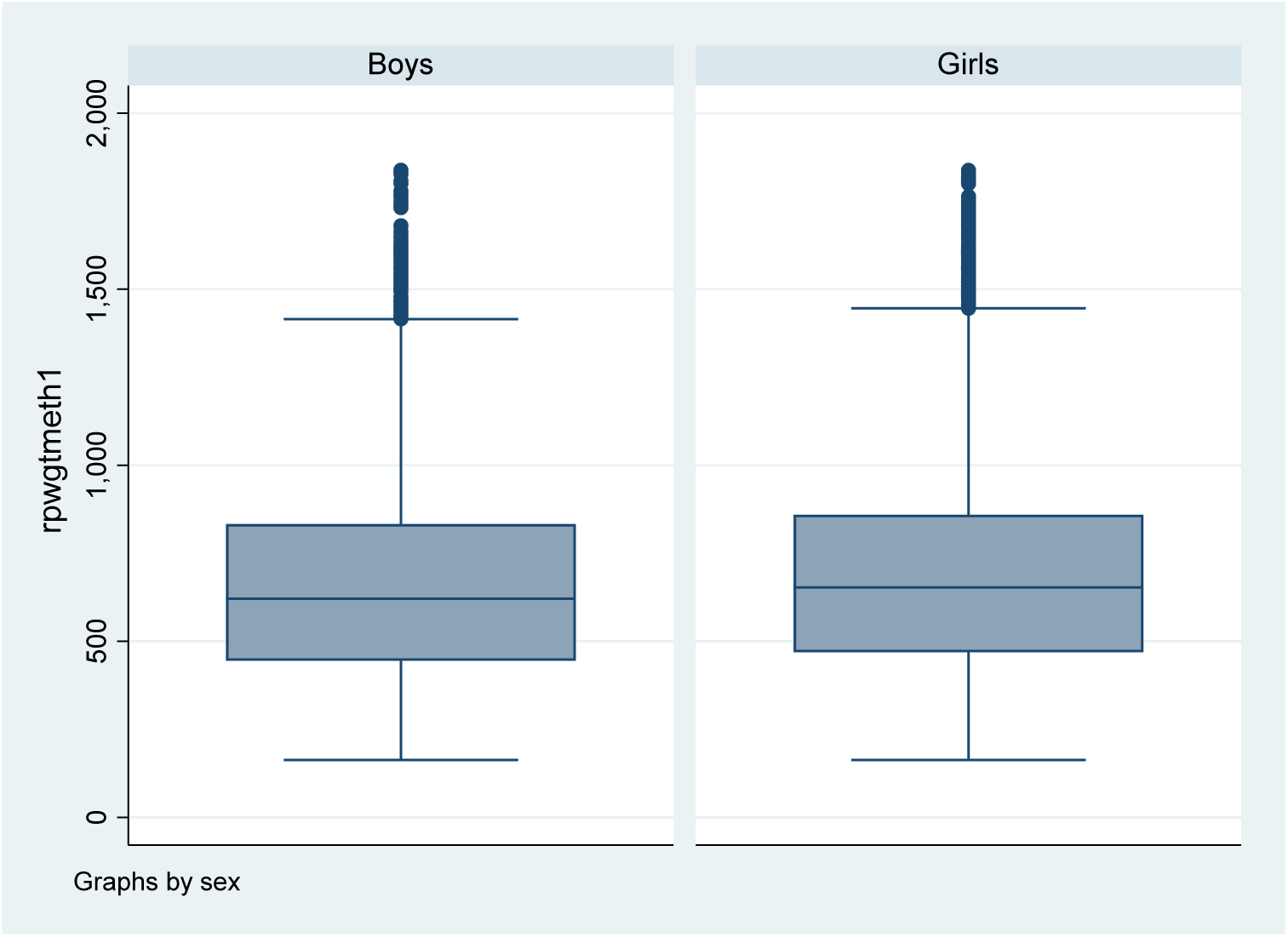
Distribution of ABCD Baseline Population Weights by Sex of Child

**Figure 3:**
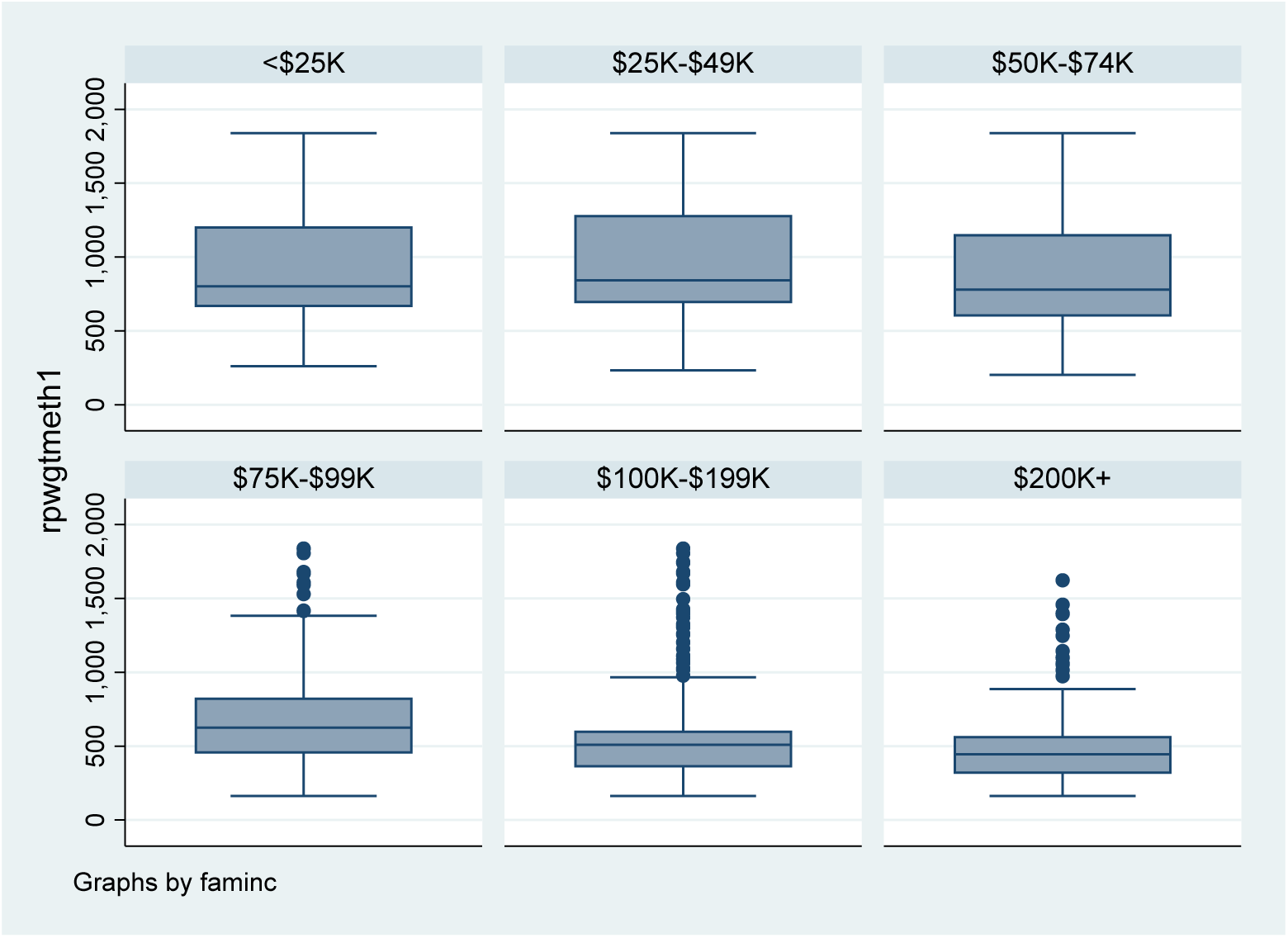
Distributions of ABCD Analysis Weights by Family Income Category

**Table 3:** U.S. Population Totals for Final Raking Step in ABCD Population Weight Calculation.

| Marginal Control Variable | Category | ACS Estimated Population |
| --- | --- | --- |
| U.s Population Age 9-10 | Total | 8,211,605 |
| Age | 9 | 4,074,807 |
|  | 10 | 4,136,798 |
| Sex | Female | 4,005,680 |
|  | Male | 4,205,925 |
| Race/Ethnicity | White | 4,305,552 |
|  | Black | 1,101,297 |
|  | Hispanic | 1,973,827 |
|  | Asian | 487,673 |
|  | All Other Children | 343,256 |

## 7. Comparison of Analysis Methods

Section 2 described a primary ABCD aim which is to enable researchers to perform analyses that represent the demographic, social, genetic and neurological characteristics of the two-year cohort of U.S. children as well as the social and environmental influences that will affect their development through their teens to young adulthood. This section compares results from a selected set of descriptive and multiple regression analyses of the ABCD baseline data conducted using the alternative design-based and model-based analysis methods described in the preceding sections.

Recognizing that no single method constitutes a gold standard, the primary aims of the comparative analyses covered in this section are to answer the following questions:

- How different are unweighted and population-weighted estimates of descriptive statistics? Are robust, design-based estimates of standard errors necessary for estimates of descriptive statistics?
- How do multi-level model estimates of regression parameters compare to quasi-probability sample (design-based) estimates for these same models.

- Do covariate controls in multi-level models achieve the same effect as propensity weights in a design-based approach?
- Are standard errors for coefficients comparable between the two approaches
- How does pooling of the ABCD general population and special twin samples impact analysis results?

- Under the design-based approach to descriptive analysis, can the two samples be pooled in a single weighted analysis or is stand-alone analysis of the general population sample preferred?
- In regression modeling, how best can the special twin sample be incorporated into multi-level modeling and design-based analyses.

The methods comparison for descriptive analyses of the population focuses on estimates and standard errors of population proportions for categorical demographic and socio-economic characteristics as well as means and distribution quantiles for continuous scale measures from the NIH Toolbox (uncorrected Flanker and Reading test scores) and counts from parental reports of the child’s lifetime visits to a hospital emergency department. Three approaches to descriptive estimation of population quantities are compared: unweighted estimation based on the full baseline sample; weighted design-based estimation for the full sample; and weighted, design-based estimation for the sample exclusive of cases in the special twin samples in four study sites.

This modest evaluation of analysis methods for the ABCD baseline data was also extended to compare various approaches to both linear and Poisson regression for selected dependent variables (outcomes) measured in the ABCD baseline assessment. The comparative investigation for linear regression of continuous dependent variables focused on models for the uncorrected Flanker and Reading test scores from the NIH Toolbox set of measures. For generalized linear regression modeling of the ABCD data, a Poisson regression model for the count of lifetime ER visits serves as the basis for comparing different methods. Six model estimation approaches were applied to each regression problem.

1. Ordinary Least Squares (OLS) or Maximum Likelihood (MLE) estimation of the model. No weighting. No design-based or model adjustments for the non-independence of observations.
2. Design-based estimation for the pooled sample. Standard statistical software for survey data analysis (R Survey Library) was applied. The pooled population weight was used along with robust (Taylor Series Linearization) variance estimation methods to account for the nested clustering of sample observations.
3. Design-based estimation for the not pooled sample. Methodology identical to (2) except that the special twin sample was excluded from the population weight development and the regression analysis.
4. Two-level (site, individual) multi-level, mixed effects model applied to the pooled sample data. Only Level 1 covariate controls for demographic and socio-economic characteristics. No population weighting used in estimating fixed effect parameters and variance components.
5. Two-level (site, individual) multi-level, mixed effects model applied to only the general population sample data (excluding special twins). Methodology identical to (4) except that the special twin sample was excluded from the regression analysis.
6. Three-level (site, family, individual) multi-level, mixed effects model applied to the pooled sample data. Only Level 1 covariate controls for demographic and socio-economic characteristics. No population weighting used in estimating Level 1 fixed effect parameters and variance components. This approximates the GAMM4 model estimation employed by default in regression analyses performed in the ABCD Data Exploration and Analysis Portal (DEAP).

### 7.A. Descriptive estimates-comparison of estimation methods

Applying the three descriptive estimation approaches described above, Table 4 compares ABCD sample estimates to the corresponding ACS demographic proportions. Due to the final raking controls employed in the process of developing the propensity-based weights, weighted estimates of age, gender and race/ethnicity proportions match the ACS control proportions exactly. Weighted estimates of proportions for family socio-economic characteristics such as family income level, family type and parental labor force status are also a close –albeit not exact— match to the ACS control values. This is in contrast to the unweighted ABCD estimates which are biased toward a population of families with married, working parents and higher income levels.

**Table 4.** ABCD Baseline Weighted and Propensity Weighted Estimates of Population Demographics

| Characteristic | Category | ACS<br>2011-2015<br>%<br>n=376,370 | ABCD<br>(Unweighted)<br>% (se)<br>n=11,874 | ABCD<br>(Weighted, Design Corrected) |  |
| --- | --- | --- | --- | --- | --- |
|  |  |  |  | Pooled<br>%(se)<br>n=11,874 | Not-Pooled<br>%(se)<br>n=9999 |
| Population | Total | 100.0 | 100.0 | 100.0 | 100.0 |
| Age | 9 | 49.6 | 52.2 (0.5) | 49.6 (1.5) | 49.7 (1.3) |
|  | 10 | 50.4 | 47.8 (0.5) | 50.4 (1.5) | 50.3 (1.3) |
| Sex | Male | 51.2 | 52.2 (0.5) | 51.2 (0.4) | 51.21 (0.45) |
|  | Female | 48.8 | 47.8 (0.5) | 48.8 (0.4) | 48.79 (0.45) |
| Race/Ethnicity | NH White | 52.4 | 52.2 (0.5) | 52.4 (5.6) | 52.4 (5.9) |
|  | NH Black | 13.4 | 15.1 (0.3) | 13.4 (2.6) | 13.4 (2.7) |
|  | Hispanic | 24.0 | 20.4 (0.4) | 24.0 (6.2) | 24.0 (6.5) |
|  | Asian, AIAN, NHPI | 5.9 | 3.2 (0.2) | 5.9 (0.5) | 5.9 (0.5) |
|  | Multiple | 4.2 | 9.2 (0.3) | 4.18 (0.6) | 4.22 (0.6) |
| Family Income | <\$25K | 21.5 | 16.3 (0.3) | 20.0 (2.4) | 19.95 (2.28) |
| | \$25K-\$49K | 21.7 | 14.9 (0.3) | 20.5 (1.6) | 20.13 (1.65) |
| | \$50K-\$74K | 17.0 | 13.8 (0.3) | 17.5 (0.9) | 16.98 (0.86) |
| | \$75K-\$99K | 12.5 | 14.3 (0.3) | 13.2 (0.8) | 13.2 (0.8) |
| | \$100K-\$199K | 20.5 | 29.6 (0.4) | 21.7 (2.2) | 22.4 (2.2) |
| | \$200K + | 6.8 | 11.2 (0.3) | 7.10 (1.0) | 7.40 (1.1) |
| Family Type | Married Parents | 66.1 | 73.5 (0.4) | 66.9 (2.1) | 67.4 (2.3) |
|  | Other Family | 33.9 | 26.5 (0.4) | 33.1 (2.1) | 32.6 (2.3) |
| Parent Employment | Married, 2 in LF | 40.8 | 50.3 (0.5) | 41.9 (1.9) | 42.2 (1.9) |
|  | Married, 1 in LF | 23.7 | 21.9 (0.4) | 23.1 (1.7) | 23.2 (1.8) |
|  | Married, 0 in LF | 1.9 | 1.3 (0.1) | 2.0 (0.3) | 1.9 (0.2) |
|  | Single, M, in LF | 26.5 | 1.6 (0.1) | 1.9 (0.2) | 2.0 (0.2) |
|  | Single, M,Not LF | 7.0 | 0.4 (0.1) | 0.4 (0.1) | 0.4 (0.1) |
|  | Single, F, in LF | xx | 19.5 (0.4) | 24.0 (1.6) | 22.4 (2.2) |
|  | Single, F,Not LF | xx | 5.1 (0.2) | 6.7 (0.6) | 7.4 (1.1) |

The comparison of estimates for these demographic and SES category proportions points to the importance of including the effects of sample clustering in the estimation of standard errors. Standard errors for the unweighted estimates that assume independence of observations seriously underestimate the design-corrected standard errors. The degree of underestimation is related to the intra-cluster correlation of the attribute and is most extreme for characteristics that are more highly clustered in a small number of sites (i.e. African American children).

The Table 4 comparative summary for demographic and socio-economic characteristics of U.S. 9 and 10 year-olds shows only minor differences in weighted estimates for the pooled and not pooled samples. This finding lends support to the study decision to employ population weighting that integrates the special twin samples with the general population sample. The obvious benefit of the pooled sample is the increase in nominal sample size that in turn results in moderately lower standard errors for total population estimates.

Table 5 extends the comparison of weighted and unweighted estimation approaches to descriptive estimation of means and population quantiles for several substantive variables measured in the ABCD baseline assessment. In this comparison, there also is little difference in results for weighted estimates based on the pooled and not pooled samples. Furthermore, unweighted and weighted estimates for the means and percentiles of the NIH Toolbox measures and lifetime ER visits exhibit at most only minor differences However, standard errors that result from the unweighted estimation are certainly underestimates of the true sampling variability of the estimates for the clustered sample observations.

**Table 5:** ABCD Baseline Estimated Distributional Statistics for NIH Tool Box Flanker and Reading Scores

| Variable | Distribution Statistic | ABCD (Unweighted) | ABCD (Weighted, Design Corrected) |  |
| --- | --- | --- | --- | --- |
|  |  |  | Pooled | Not-Pooled |
| NIH Toolbox Flanker Test (Uncorrected) | n | 11,712 | 11712 | 9999 |
|  | Mean | 94.0 (0.08) | 93.83 (0.30) | 93.83 (0.27) |
|  | Q5 | 75.98 (0.34) | 75.52 (0.96) | 75.62 (0.99) |
|  | Q25 | 88.95 (0.14) | 88.70 (0.50) | 88.72 (0.48) |
|  | Q50 (Median) | 94.94 (0.10) | 94.79 (0.26) | 94.76 (0.22) |
|  | Q75 | 99.79 (0.09) | 99.65 (0.19) | 99.64 (0.16) |
|  | Q95 | 105.77 (0.11) | 105.74 (0.18) | 105.77 (0.16) |
| NIH Toolbox Reading Test (Uncorrected) | n | 11704 | 11704 | 9991 |
|  | Mean | 90.86 (0.06) | 90.60 (0.07) | 90.74 (0.25) |
|  | Q5 | 79.22 (0.18) | 78.77 (0.49) | 78.71 (0.48) |
|  | Q25 | 88.69 (0.09) | 86.40 (0.28) | 86.46 (0.29) |
|  | Q50 (Median) | 90.25 (0.04) | 90.05 (0.10) | 90.20 (0.11) |
|  | Q75 | 94.26 (0.06) | 94.00 (0.31) | 94.25 (0.16) |
|  | Q95 | 101.37 (0.12) | 101.25 (0.25) | 101.47 (0.25) |

### 7.B Linear Regression analyses—comparison of model fitting approaches

Tables 6 and 7 summarize results from the various approaches to fitting the linear regression models for the two NIH Toolbox measures. Table 6 presents the full set of parameter estimates and standard errors for the OLS fit of the models for the uncorrected Flanker and Reading Test Scores. Focusing substantively on the main effects of age and income for these Toolbox scores, the OLS estimates suggest that, all else being equal, age 9 children have uncorrected Flanker Scores and Reading Scores that are approximately 2.7 units lower than their 10 year old counterparts. After controlling for other main effects, children from the highest income families ($200K+) average 4.5-5.5 points higher on the tests compared to children from the lowest income families.

**Table 6:**
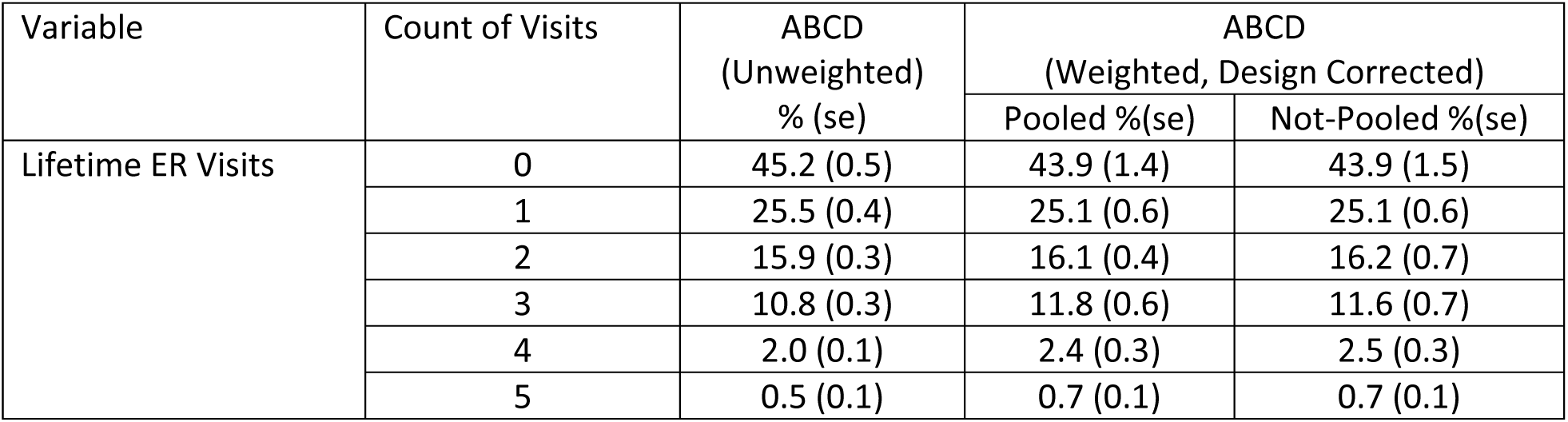
Estimates of Child’s Lifetime Visits to Emergency Room (ER)

**Table 6:** ABCD Baseline OLS Regression of NIH Toolbox Flanker and Reading Test Scores

| OLS Regression Parameter | Flanker (Uncorrected) |  |  |  | Reading (Uncorrected) |  |  |  |
| --- | --- | --- | --- | --- | --- | --- | --- | --- |
| | Est($\beta$) | Se($\beta$) | CIL($\beta$) | CIU($\beta$) | Est($\beta$) | Se($\beta$) | CIL($\beta$) | CIU($\beta$) |
| Intercept | 93.75 | 0.50 | 92.77 | 94.73 | 86.83 | 0.34 | 86.16 | 87.50 |
| Age:9 | **-2.69** | **0.16** | **-3.00** | **-2.38** | **-2.65** | **0.12** | **-2.89** | **-2.41** |
| Sex:Female | -0.30 | 0.16 | -0.61 | 0.01 | 0.08 | 0.12 | -0.16 | 0.32 |
| RaceEth:Asian/Oth | 1.50 | 0.47 | 0.58 | 2.42 | 1.11 | 0.36 | 0.40 | 1.82 |
| Race\Eth:Hispanic | -0.37 | 0.23 | -0.82 | 0.08 | -1.00 | 0.17 | -1.33 | -0.67 |
| RaceEth: Multiple | 0.34 | 0.29 | -0.23 | 0.91 | 0.10 | 0.22 | -0.33 | 0.53 |
| RaceEth:Black | -2.74 | 0.31 | -3.35 | -2.13 | -2.56 | 0.20 | -2.95 | -2.17 |
| Family_Employment: Married, 0 in LF | -0.87 | 0.82 | -2.48 | 0.74 | 1.00 | 0.60 | -0.18 | 2.18 |
| Family_Employment: Married, 1 in LF | -0.54 | 0.47 | -1.46 | 0.38 | 1.55 | 0.32 | 0.92 | 2.18 |
| Family_Employment: Married,2 in LF | -0.49 | 0.46 | -1.39 | 0.41 | 1.48 | 0.31 | 0.87 | 2.09 |
| Family_Employment: Single HH, 1LF | -0.17 | 0.46 | -1.07 | 0.73 | 1.41 | 0.31 | 0.80 | 2.02 |
| Faminc :25k-49k | 1.70 | 0.34 | 1.03 | 2.37 | 1.82 | 0.23 | 1.37 | 2.27 |
| Faminc: 50k-74k | 2.74 | 0.36 | 2.03 | 3.45 | 3.37 | 0.24 | 2.90 | 3.84 |
| Faminc: 75k-99k | 3.17 | 0.36 | 2.46 | 3.88 | 3.75 | 0.25 | 3.26 | 4.24 |
| Faminc: 100k-199k | 3.78 | 0.34 | 3.11 | 4.45 | 4.45 | 0.23 | 4.00 | 4.90 |
| Faminc: 200k+ | **4.51** | **0.39** | **3.75** | **5.27** | **5.57** | **0.27** | **5.04** | **6.10** |
| sitetwin 0 | -0.10 | 0.22 | -0.53 | 0.33 | 1.53 | 0.16 | 1.22 | 1.84 |

**Table 7:**
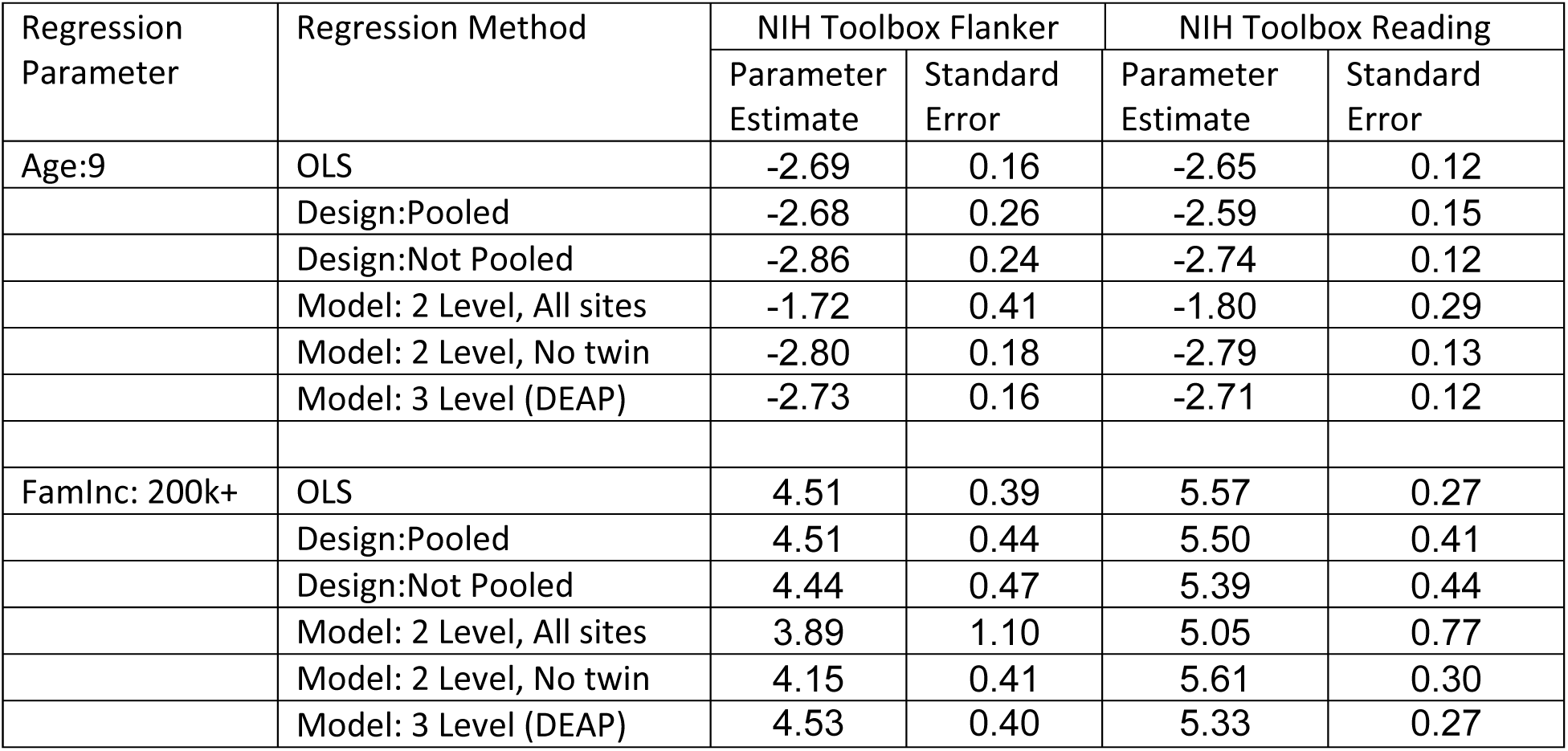
Comparison of selected linear regression parameters. NIH Toolbox Uncorrected Flanker and Reading Test. Source: ABCD Baseline.

Table 7 selects these two important main effects from the OLS model fit (AGE, INCOME LEVEL>$200K) for the Toolbox test scores and compares the estimated parameters and standard errors that result when the six different design-based and model-based estimation methods are applied. Although a true gold standard is lacking and it is important not to over generalize, the following paragraphs compare the OLS and Linear Mixed Model (LMM) fits of the regressions followed by a comparison of the two and three-level mixed effects models. Results for the three-level mixed effects model-based estimates are then compared to weighted design-based estimates.

#### 7.B.1 OLS vs. Linear Mixed Effects Models (LMM)

The OLS and three-level LMM approaches produce very similar estimates of the fixed effects of age and income level on the Flanker and Reading test scores regressands. Likewise, the OLS and LMM estimates of the standard errors of the estimated coefficients for these two regression models are very similar.

#### 7.B.2 LMMs for ABCD: Three vs. Two Levels?

The comparative results for these two regression models suggest that when the special ABCD twin sample data are pooled with the general population sample and a LMM approach is used it is important to apply the three level DEAP model that includes a level two contribution for clustering within family unit. When a two level model is applied to these pooled data and family level clustering is ignored, the results of these example analyses suggest the parameter estimates will be attenuated and estimated standard errors will be seriously overestimated. If the two level LMM is fitted using only data for the general population sample (excluding the special twin sample cases), the resulting parameters estimates and standard errors are more consistent with those for the three level model.

#### 7.B.3 Three-level LMM vs. Design-based Population weighted LS and Robust SEs

As noted above, the unweighted LMM and weighted design-based approaches compared in Table 7 aim to capture/model the complex variance structure of clustering and non-independence of the baseline observations for the ABCD child cohort. The design-based estimation approaches employ the population weights described in Section 6 above and use a weighted least squares (WLS) methodology to estimate the population regression parameters. Under the design-based approaches, a Taylor Series Approximation (or sandwich estimator) is used to compute robust estimates of standard errors. However, unlike the LMM approach, the components of variance associated with each level of clustering are estimated as a single weighted aggregate for the residual variance and not as individual components of variance attributable to each level of the clustering. The three-level LMM used here does not include population weighting in estimating the regression parameters. The three-level LMM does produce estimates of the variance components associated with random site effects, family effects and the residual variance for the individual child observations.

The three-level LMM and design-based approaches produce very similar estimates of the age and income parameters for the Flanker test regression model; however, estimated standard errors are higher under the designed based model fit—a loss in precision that may be attributed to the variability (see Figure 1) in population weights and a weak correlation between those weights and the Flanker score itself. Switching to the Reading test regression, the comparison is more mixed, particularly for estimates of the income parameter. The three-level LMM estimate for the income coefficient is considerably lower than the design-based estimate based on the pooled data. If the uncertainty of estimates is taken into account this difference is not significant. The difference in the estimated effect of high income may be attributed to the design-based weighting which as illustrated in Figure 3 varies substantially as a function of family income level.

### 7.C Generalized Linear Model: Poisson Regression—Comparison of model fitting methods

Table 8 presents a summary of standard, unweighted maximum likelihood estimates of parameters for the Poisson regression of the count of children’s lifetime ER visits (e.g. 0, 1,2,…) on the selected set of demographic and socio-economic covariates. Focusing again on the specific effect of age (9 vs. 10) and family income ($25-49K vs <$25K), Table 8 compares various analytic approaches to estimating the relative risk for the younger children and those from higher income families. Here again, as in the previous comparisons based on the linear regression model, the three-level DEAP LMM and the design-based estimation for the pooled data show minor differences in the estimated relative risks and confidence intervals but the magnitude of these differences would not be judged to be substantively important. The largest difference between estimates produced by these two methods occurs in estimation of the income effect which again is likely attributable to the weighting adjustment that is included in the design-based estimates.

**Table 8:** Poisson Regression of Lifetime ER Visit Counts. Source: ABCD Baseline.

| MLE<br>Parameter | Lifetime Visits to ER |  |  |  |  |
| --- | --- | --- | --- | --- | --- |
|  | Model Coefficients |  | Relative Risk |  |  |
|  | Parameter | Standard Error | Estimate | LCI | UCI |
| **Intercept** | 0.303 | 0.050 | - |  | - |
| **Age:9** | -0.034 | 0.018 | 0.97 | 0.93 | 1.00 |
| **Sex:Female** | -0.148 | 0.0186 | 0.86 | 0.83 | 0.89 |
| **RaceEth:Asian/Oth** | -0.31 | 0.063 | 0.73 | 0.65 | 0.83 |
| **Race\Eth:Hispanic** | -0.039 | 0.026 | 0.96 | 0.91 | 1.01 |
| **RaceEth: Multiple** | 0.104 | 0.02 | 1.11 | 1.07 | 1.15 |
| **RaceEth:Black** | -0.127 | 0.031 | 0.88 | 0.83 | 0.94 |
| **Family_Employment: Married, 0 in LF** | -0.316 | 0.093 | 0.73 | 0.61 | 0.87 |
| **Family_Employment: Married, 1 in LF** | -0.228 | 0.044 | 0.80 | 0.73 | 0.87 |
| **Family_Employment: Married,2 in LF** | -0.212 | 0.043 | 0.81 | 0.74 | 0.88 |
| **Family_Employment: Single HH, 1LF** | -0.064 | 0.042 | 0.94 | 0.86 | 1.02 |
| **Faminc :25k-49k** | 0.131 | 0.033 | 1.14 | 1.07 | 1.22 |
| **Faminc: 50k-74k** | 0.1 | 0.036 | 1.11 | 1.03 | 1.19 |
| **Faminc: 75k-99k** | -0.062 | 0.038 | 0.94 | 0.87 | 1.01 |
| **Faminc: 100k-199k** | -0.079 | 0.036 | 0.92 | 0.86 | 0.99 |
| **Faminc: 200k+** | -0.111 | 0.043 | 0.89 | 0.82 | 0.97 |
| **sitetwin 0** | -0.011 | 0.027 | 0.99 | 0.94 | 1.04 |

**Table 9:** Poisson Regression for Count of Lifetime ER Visits. Source: ABCD Baseline.

| Regression Parameter | Regression Method | Model Coefficient |  | Relative Risk Ratio |  |  |
| --- | --- | --- | --- | --- | --- | --- |
|  |  | Parameter Estimate | Standard Error | Relative Risk | LCI | UCI |
| Sex: Female | MLE | -0.148 | 0.019 | 0.86 | 0.83 | 0.90 |
|  | Design:Pooled | -0.135 | 0.020 | 0.87 | 0.84 | 0.91 |
|  | Design:Not Pooled | -0.130 | 0.020 | 0.88 | 0.84 | 0.91 |
|  | Model: 2 Level, All sites | -0.149 | 0.016 | 0.86 | 0.83 | 0.89 |
|  | Model: 2 Level, No twin | -0.151 | 0.016 | 0.86 | 0.83 | 0.89 |
|  | Model: 3 Level (DEAP) | -0.145 | 0.021 | 0.87 | 0.83 | 0.90 |
| FamInc: 25-49k | MLE | 0.131 | 0.033 | 1.14 | 1.07 | 1.22 |
|  | Design:Pooled | 0.113 | 0.041 | 1.12 | 1.03 | 1.21 |
|  | Design:Not Pooled | 0.120 | 0.045 | 1.13 | 1.03 | 1.23 |
|  | Model: 2 Level, All sites | 0.144 | 0.039 | 1.15 | 1.07 | 1.25 |
|  | Model: 2 Level, No twin | 0.152 | 0.039 | 1.16 | 1.08 | 1.26 |
|  | Model: 3 Level (DEAP) | 0.141 | 0.040 | 1.15 | 1.06 | 1.25 |

## 8. Summary: Recommendations for research analysts

Although it is important not to over generalize from a small set of comparative analyses to all possible analyses of the ABCD data, the results described in section 8 lead to several recommendations for researchers who are analyzing the ABCD baseline data.

1. Unweighted analysis may result in biased estimates of descriptive population statistics. The potential for bias in unweighted estimates from the ABCD data is strongest when the variable of interest is highly correlated with socio-economic variables including family income, family type and parental work force participation. Robust variance estimation methods for clustered sample data should be employed in order not to underestimate the true uncertainty associated with estimates of descriptive statistics. For descriptive estimates of population characteristics, attitudes and behaviors and measures of estimate uncertainty including standard errors and confidence intervals, the recommend approach is to employ robust, population weighted, design based estimation using software developed for analysis of complex sample survey data. The ABCD DEAP employs the R Survey Package (Lumley, 2010) as the default for producing descriptive estimates for the population of ABCD children. Analysts interested in stand-alone analysis of the ABCD baseline data should employ the programs in the R Survey Library or survey-based procedures available in one of the major software packages listed in attachment A. In specifying the design parameters for these programs, ABCD site would be declared as the ultimate cluster (UC) or primary sampling unit (PSU) variable and the propensity-based population weight as the analysis weight variable.
2. For regression modeling of the ABCD baseline using DEAP, the three-level (site, family, individual) multi-level specification is the preferred choice. Regression analyses conducted using DEAP will default to fitting the model using the R system GAMM4 package with the three-level nested clustering specification for sites, family units and individuals. Presently, there is no empirical evidence from preliminary comparative analysis trials (results not reported here) that methods for multi-level weighting (Rabe-Hesketh and Skrondal, 2006) will improve the accuracy or precision of the model fit although additional research on this topic is ongoing.
3. As an alternative to the DEAP multi-level modeling, analysts conducting regression analysis of the ABCD baseline data outside of the NDA/DEAP environment can choose to employ the R system GAMM4 package directly or employ multi-level modeling programs available in other major software packages such as SAS, Stata, or MPlus. Following the default method employed in DEAP, the basic multi-level model should be specified to include random effects at Level 2 (family) and Level 3 (site). Analysts may also opt to use standard survey software packages (see Appendix A) and conduct design-based estimation of the regression relationship. Under the design based approach, the ABCD site would be declared as the UC or PSU variable and the propensity-based population weight should be used. Appendix B provides example syntax for specifying the ABCD design variables (cluster code, propensity weight) in the survey commands in the SAS®, STATA® and R statistical software systems.
4. The comparative analyses of descriptive estimation methods presented in Section 7.A found that, properly weighted, results for the pooled general population and special twin samples are comparable to those for weighted estimates based solely on the smaller general population sample. Likewise, for the selected models, regression analyses based on the pooled general population and special twin samples that account for inter-familial clustering (either the DEAP three-level model or design-based model estimation) produce similar results to analyses based on the general population sample alone. Nevertheless, analysts should use appropriate caution in pooling the general population and special twin samples for all forms of ABCD analysis. After controlling for demographic and socio-economic effects, the outcomes examined here—test scores, ER visits—do not appear to different substantially for twins and single births. Such exchangeability may not necessarily hold in other ABCD analyses (e.g. investigation of genetic effects or familial/sibling influences on development).
5. Finally, this guide has addressed many of the sources of statistical uncertainty faced by analysts of the ABCD baseline data. The comparison of analysis methods and general recommendations reported here serve as a starting point for analyses of these data. Researchers are encouraged to consider each of the informative features of the ABCD (clustering, sample selectivity, twin sample pooling) as they may apply to their analytic aims. Sensitivity analyses such as those underlying the comparisons in Section 7 should provide good insight into the degree to which results for descriptive estimates or fitted models are influenced by clustering, weighting and twin sample pooling.

## Appendix A: Software for Analysis of Survey Data

- SAS® software, Proc SurveyXXXX http://www.sas.com
- STATA® software, surveyset and svy: modifier http://www.stata.com
- R software, Survey library (Lumley, 2010) http://www.r-project.org/
- Sudaan® software http://www.rti.org
- SPSS® software, Complex samples module http://www.spss.com
- Mplus® software http://statmodel.com
- WesVar http://www.westat.com/westat/statistical_software/wesvar
- IVEware http://www.isr.umich.edu/src/smp/ive
- DEAP – https://scicrunch.org/resolver/SCR_016158 Library

## Appendix B: Example Syntax for Analysis of ABCD Data Using Survey Analysis Software

### SAS V9+

**proc surveymeans** data=abcd2.abcd_092017_test;

weight rpwgtmeth1;

cluster abcd_site;

var nihtbx_flanker_uncorrected nihtbx_reading_uncorrected;

**run**;

**proc surveyreg** data=abcd2.abcd_092017_test;

class ageyr sex c_race_eth c_fesabcd faminc sitetwin;

weight rpwgtmeth1;

cluster abcd_site;

model nihtbx_flanker_uncorrected=ageyr sex c_race_eth c_fesabcd faminc sitetwin /solution;

**run**;

### STATA

**svyset abcd_site** [**pweight=rpwgtmeth1**]

svy:mean nihtbx_flanker_uncorrected nihtbx_reading_uncorrected svy: regress nihtbx_flanker_uncorrected i.ageyr i.sex i.c_race_eth /// i.c_fesabcd i.faminc i.sitetwin

### R Survey Package

\# Survey package library(survey)

\# set survey data

svyr <-svydesign(data=statadata, id=∼site_name, strata=NULL, weights=statadata$rpwgtmeth1)

\# weighted w/complex sample design correction

svymean(∼nihtbx_flanker_uncorrected, design=svyr, na.rm=T) svymean(∼nihtbx_reading_uncorrected, design=svyr, na.rm=T)

\# with linear outcome of Flanker with design correction

summary(reg_ex1 <-svyglm(nihtbx_flanker_uncorrected ∼ agef + sexf + c_race_ethc + fesf + famincc + sitetwinc, design=svyr))

